# Encoding of speech in convolutional layers and the brain stem based on language experience

**DOI:** 10.1101/2022.01.03.474864

**Authors:** Gašper Beguš, Alan Zhou, T. Christina Zhao

**Affiliations:** Department of Linguistics, University of California, Berkeley, United States; Department of Cognitive Science, Johns Hopkins University, United States; Institute for Learning and Brain Sciences, University of Washington, United States; Department of Speech and Hearing Sciences, University of Washington, United States

## Abstract

Comparing artificial neural networks with outputs of neuroimaging techniques has recently seen substantial advances in (computer) vision and text-based language models. Here, we propose a framework to compare biological and artificial neural computations of spoken language representations and propose several new challenges to this paradigm. The proposed technique is based on a similar principle that underlies electroencephalography (EEG): averaging of neural (artificial or biological) activity across neurons in the time domain, and allows to compare encoding of any acoustic property in the brain and in intermediate convolutional layers of an artificial neural network. Our approach allows a direct comparison of responses to a phonetic property in the brain and in deep neural networks that requires no linear transformations between the signals. We argue that the brain stem response (cABR) and the response in intermediate convolutional layers to the exact same stimulus are highly similar and quantify this observation. The proposed technique not only reveals similarties, but also allows for analysis of the encoding of actual acoustic properties in the two signals: we compare peak latency (i) in cABR relative to the stimulus in the brain stem and in (ii) intermediate convolutional layers relative to the input/output in deep convolutional networks. We also examine and compare the effect of prior language exposure on the peak latency in cABR and in intermediate convolutional layers. Substantial similarities in peak latency encoding between the human brain and intermediate convolutional networks emerge based on results from eight trained networks (including a replication experiment). The proposed technique can be used to compare encoding between the human brain and intermediate convolutional layers for any acoustic property and for other neuroimaging techniques.

## 1 Introduction

Though several characteristics of artificial neural networks (ANNs) are biologically implausible, many aspects of ANNs are biologically inspired and have equivalents in the human brain^1–4^. Among the architectures highly influenced by brain processing in the visual domain are convolutional neural networks (CNNs)^5–10^. Despite the fact that many aspects of current ANNs lack biological plausibility, it is still reasonable to compare computations and representations in deep neural networks and the brain. Such work has twofold implications. On the one hand, the comparison has the potential to shed light on how ANNs encode representations internally relative to the brain and how learning biases in humans and ANNs differ. On the other hand, computational models allow us to simulate brain processes (such as speech) and test hypotheses that are not possible to test in the human brain. Such simulations can bring insights for how language gets acquired and encoded in the brain. For example, we can test what properties of speech (both in terms of behavioral and neural data) emerge when models have no articulatory biases compared to models with articulatory information, or when models have no language-specific mechanisms compared to models with language-specific biases. Such simulations can help us better understand which properties of language are domain specific vs. domain general, and which properties emerge from articulatory or cognitive factors (Section 1.3).

The majority of work comparing the brain and ANNs is performed on the visual domain, with substantially less work comparing ANNs to brain responses to linguistic stimuli. Most existing comparison studies in the linguistic domain focus on text-trained models and supervised models, and focus on correlations. Here, we outline a technique that parallels biological and artificial neural encoding of specific acoustic phonetic features by analyzing ANN models trained on raw speech in a fully unsupervised manner. We introduce the GAN architecture^11^ to the brain-ANN comparison literature.

GANs are uniquely appropriate for modeling speech acquisition^12,13^. Crucially, GANs need to learn to generate output from noise by imitation/imagination in a fully unsupervised manner. The main characteristic of the architecture are two networks, the Generator and the Discriminator, that are trained in a minimax game^11^, in which the Discriminator attempts to distinguish real data and outputs from the Generator, and the Generator learns to generate realistic outputs given only feedback from the Discriminator (summary in Figure 2). It has been shown that this process results in the ability to encode linguistic information (e.g. lexical and sublexical representations) into raw speech in a fully unsupervised manner^13^ as well as in the ability to learn highly complex morphophonological rules^14^ both locally and non-locally^15^. In other words, linguistically meaningful representations (such as words, prefixes) self-emerge in the GAN architecture when the models are trained on raw speech. Evidence for several hallmarks of symbolic-like representations emerges in GANs: discretized (disentangled) representations, a causal relationship between the latent space and generated outputs, and near-categoricity of desired outputs^13,14^. Crucially, GANs do not simply replicate input data or predict the next sequence (as is the case for other deep learning models such as autoencoders or text-based transformers), but generate innovative and interpretable outputs from noise, which mimics one of the more prominent features of language: productivity^16^.

### 1.1 Prior work

A substantial amount of work exists on paralleling brain imaging with artificial neural networks in the visual domain^7,9,17–23^ and relatively fewer works exist in the language or in the speech domain (most works focus on text-based language models^24–26^). Kell et al.^27^ parallel fMRI recordings with supervised speech and music recognition model trained on waveforms, while Millet and King^28^ parallel fMRI recordings with ASR models trained on spectrograms. The comparisons^27,28^ reveal parallels in neural encoding between ANNs and the brain, but are based on a linear regression estimates between the two sets of signals. They also focus on correlations, and do not directly compare individual acoustic properties without linear transformations. While the approach to ANN-brain comparison that uses linear transformations between the signals can operate with more complex data (such as in Kell et al.^27^), transformation also decreases the interpretability of the comparison. Huang et al.^29^ examine a measurement of surprisal in a supervised (CNN) classifier, and correlate the metric to an EEG signal reduced in dimensionality. Donhauser and Baillet^30^ train a predictive ANN model and use it to quantify the brain’s response to surprisal during speech processing. Koumura et al.^31^ focus on amplitude modulation of auditory stimuli (not only of speech). Their model is trained on raw waveforms, but the analysis focuses on individual units in deep convolutional networks. They analyze synchrony and average activity for each unit and analyze them across convolutional layers. All their models are fully supervised classifiers (thus modeling only perception) and do not focus on linguistically meaningful representations, but on acoustic phonetic properties of speech and audition in general. Smith et al.^32,33^ argue for parallels in human binaural detection and deep neural networks (variational autoencoders or VAEs). They model pure tones rather than speech and focus on binaural detection. Khatami and Escabí^34^ operate with hierarchical spiking neural networks on cochleograms using supervised training and parallel the resulting model with the hierarchical organization of the human auditory system. Magnuson et al.^35^ compare a classifier (based on *long short-term memory* or LSTM) trained on spectrograms to electrocorticography (ECoG) data. Most of these proposals focus on correlations, similarity scores, or linear transformations between ANN and brain representations. The speech datasets in all studies except in Millet and King^28^ are limited to one language—English from TIMIT or from other corpora. Saddler et al.^36^ compare supervised deep convolutional networks for F0 classification with models of the auditory nerve, but not with actual brain imaging data. All these frameworks use supervised classification networks for their comparison. Below, we outline how our model differs from these existing proposals.

### 1.2 Goals & new challenges

This paper proposes some crucial new approaches and guidelines to the comparison of how deep neural networks and the brain represent spoken language. First, we compare brain data to fully unsupervised models where linguistically meaningful representations need to self-emerge. Language acquisition is predominantly unsupervised with only some limited aspects of acquisition being implicitly or explicitly supervised (such as negative feedback^37,38^). Rather than analyzing pre-trained models, we also custom-train the networks on controlled data which allows for more interpretable results and a more direct comparison with human experiments. For example, we can train the network on the same speech process that is tested in the brain-imaging experiment (such as aspiration of stops) or test encoding in ANNs using the exact same stimulus that is tested in brain-imaging experiment. Smaller training datasets also more closely resemble language acquisition in initial stages when the number of lexical items is highly limited^39^.

Second, our models and visualization techniques capture both the production and perception component in human speech (equivalent to the encoding and decoding, two central concepts in cognitive science^40^), while most existing proposals exclusively focus on the perception component. We perform comparison between brain and ANN data from both the Generator network that simulates speech production (synthesis, decoding) and the Discriminator network that simulates speech perception (classification, encoding). For modeling the production element, we propose a procedure for comparing ANNs with the brain data where the model’s internal elements (latent space) are chosen such that the model’s generated output and the stimulus in the neuroimaging experiment are maximally similar (Section 4.2; to force similarity we use a techniques in Lipton and

Tripathi^41^ and Keyes et al.^42^).^1^ For modeling the perception element, we feed the Discriminator network the actual stimulus (Section 4.3) as well as the outputs of the Generator that are forced to resemble the stimulus (Section 4.4). The production and perception in human speech are highly interconnected^44^, which is why modeling both principles is desired when comparing brains and ANNs.

Third, instead of focusing on correlations or linear transformations between signals in neuroimaging experiments and values of internal layers in deep neural networks, we focus on comparing actual acoustic features across the two systems directly, with no transformations or correlations. We argue that the two signals are highly similar even without any transformations. We analyze peak latency in both the cABR and in deep convolutional neural networks. This is a measurable acoustic property, is directly comparable, and requires no computation of correlations or any linear transformations/regressions between signals. Comparing acoustic properties rather than correlations is more interpretable: correlations can arise even in untrained models and are generally problematic to analyze and interpret.

Fourth, most of the existing proposals focus on correlating brain responses and outputs of neural networks in a single language. Monolingual comparisons primarily model acoustic encoding of speech signal and do not provide information on encoding of phonological contrasts across languages. By training the networks on two languages with a different encoding of a phonetic property (as confirmed by brain experiments), we not only test the encoding of acoustic properties, but also of phonetic features that constitute phonological contrasts: the distinction between voiceless stops (such as [t]) and voiced stops (such as [d]) in English and Spanish. Probing how *phonological* (meaning-distinguishing) contrasts are encoded in the brain and in deep neural network trained on speech can yield new information on encoding of linguistically meaningful units across the two systems.

Fifth, we propose a technique to compare EEG signals to intermediate representations in deep neural networks (for a comparison between EEG signals and ANNs in the visual domain, see Greene and Hansen^21^; for speech, see Huang et al.^29^). Unlike other neuroimaging techniques (e.g. fMRI or ECoG), EEG is minimally invasive while providing high temporal resolution, which is crucial for examining temporally dynamic speech encoding. This should allow a large-scale comparison between deep neural networks and the brain not only for those phonetic properties investigated in this paper, but for any other acoustic property.

Finally, we argue that earlier layers in deep neural networks correspond to earlier stages of speech processing in the brain. For this reason, we focus on the complex auditory brainstem response (cABR), a potential that can robustly reflect sensory encoding of auditory signals in early stages of auditory processing^45^. Comparing cABRs and deep networks is, to our knowledge, new in the paradigm of comparing deep learning and the brain. Unlike other imaging techniques (such as fMRI or ECoG), cABR is one of the few brain imaging techniques that allows recording of the brainstem regions and captures the earliest stages of speech processing. Recent evidence suggests that several acoustic properties that result in phonological contrasts are encoded already in the brain stem^46,47^.

To achieve these goals, we compare outputs of the cABR experiment^46^ to ANN representations in intermediate layers closest to the stimulus (the fourth/first convolutional layer out of five total layers) in the production/perception network, respectively. The networks are trained in a Generative Adversarial Network framework^11^, where the Generator network learns to produce speech from some random latent distribution and the Discriminator learns to distinguish real from generated samples. In other words, the Generator needs to learn to produce speech-like units in a fully unsupervised way — it never actually accesses real data, but rather needs to trick another network by producing real-looking data outputs. This unsupervised learning process based on imitation/imagination, where the networks learn to generate data from noise based only on unlabeled data, closely resembles language acquisition^12^. We train the networks on sound sequences that are acoustically similar to the stimulus in the cABR experiments and are sliced from two corpora — one on English (TIMIT^48^) and one on Spanish (DIMEx^49^), simulating the monolingual English and Spanish subjects in the cABR experiment.

We propose a new technique for comparing neuroimaging data and outputs of deep neural networks. To analyze internal representations of the network that simulates production of speech, we force the Generator to output sounds that closely resemble the stimulus used in the cABR experiment. To analyze internal representations of the network that simulates perception of speech, we feed these generated outputs as well as the actual stimulus used in the brain experiment to the Discriminator network. Using the visualization techniques proposed in Beguš and Zhou^50,51^, we can analyze any acoustic property of speech in internal convolutional layers in either the Generator (simulating speech production) or the Discriminator network (simulating speech perception). The comparison is then performed between (i) the generated outputs/stimulus in deep neural networks, and corresponding values in the second-to-last convolutional layer in the Generator/the first convolutional layer in the Discriminator and (ii) the stimulus played to subjects during the experiment and averaged cABR recording in the brain stem. We argue that this technique yields interpretable results — we can take any acoustic property with frequencies below the limit for cABRs and compare its encoding in the brain and in the artificial neural networks. To test how language experience alters representations in the brain and in artificial neural networks, we perform the comparisons on monolingual subjects of two languages in the neuroimaging experiment and deep learning models trained on the same two languages.

The results in this paper suggest that brain stem (cABR) responses and responses in the intermediate convolutional layers to to the exact same stimulus are highly similar and that peak latency differs in similar ways in the brain stem and in deep convolutional neural networks depending on which languages subjects/models are exposed to. To avoid idiosyncrasies in the models, we replicate the experiment and test encoding of both the actual stimulus and generated data. Results are consistent across sets of generated outputs and averaged stimulus inputs from four independently trained models.

### 1.3 Limitations of comparison between brains and deep neural networks

Comparing representations and computations in the human brain and deep learning models is a complex task. The goal of this paper is not to argue that human speech processing operates exactly as in deep convolutional networks (for a general discussion, see Guest and Martin^52^). We do, however, argue that computations and encodings are similar in interpretable ways between the two signals and that they result from similar underlying mechanisms (Section 5.2). These similarities set the basis for further modeling work that has the potential to offer insights both into how humans acquire and process speech as well as into how deep learning models learn internal representations.

For example, our models are closer to reality than most existing models because the learning is fully unsupervised, the models are trained on raw speech which requires no preabstraction or feature extraction^12,14^, and the CNN architecture is biologically inspired and in many ways realistic^53,54^. The models, however, still feature several unrealistic properties (beside backpropagation^54^). First, our models are trained exclusively on adult directed speech and do not include any visual information. While most models including ours disregard the visual component in language acquisition, unsupervised models still resemble human speech acquisition more closely than supervised models trained for automatic speech recognition or acoustic scene classification tasks. Additionally, we train the networks on a subset of syllables that are possible in English and Spanish (Section 3.2).

Second, we use one-dimensional CNNs for the ANN-brain comparison because of their high temporal resolution. Other architectures that better capture the temporal aspect of speech processing (such as recurrent neural networks like LSTMs would require windowing and thus likely lose the very high temporal resolution required for the short peak latency differences observed in the brain, especially if spectral transformations are required). While CNNs lack a sequential structure, they have been shown to replicate temporal effects in speech (such as locality preference^15^).

Finally, the models do not operate directly with articulatory data (they do not generate representations of the vocal tract, but rather acoustic data), while humans acquire the ability to produce speech with articulators. While these limitations are undesired because they make models less realistic, they can also be advantageous from a cognitive modeling perspective. A long-standing debate in linguistics and speech science concerns whether typological tendencies in speech patterns across the world’s languages result from articulatory pressures and transmission of language in space and time, or from cognitive biases^55–60^. Another major debate in linguistics assesses which properties of language are domain-specific and innate and which can be explained by domain-general cognitive principles^61^. Modeling speech processing in deep neural networks that contain no articulatory representations and no language-specific elements allow us to test which linguistically meaningful representations can emerge even if the models lack these properties. Such modeling can help us understand how human language is affected by cognitive, domain-general, and articulatory pressures. Combining the technique proposed in this paper with the ArticulationGAN model^62^ that introduces articulatory representations to GANs, will additionally allow us to test how articulation influences learning of linguistic representations not only behaviorally, but also with respect to artificial and biological neural computation.

## 2 cABR Experiment

The complex auditory brainstem response (cABR) reflects the early sensory encoding of complex sounds along the auditory pathway (Figure 1b) and can be measured with a 3-electrode setup using EEG^45^. The cABR generally contains an onset component, corresponding to transient changes in acoustics (e.g. stop consonant) as well as a frequency-following-response component (FFR), corresponding to periodic portions of the sound (e.g. tone, vowel). In recent decades, there has been a growing literature on characteristics of cABR. Few studies that focused on speech perception have demonstrated evidence in support for important speech perception phenomenon at the cABR level. For example, native Mandarin speakers demonstrated FFR that tracks the pitch of the lexical tones better than English speakers, demonstrating that the language experiential effect can be observed at the encoding stage^63^. The directional asymmetry phenomenon in speech perception was also observed in FFR to vowels^47^. Further, the cABR and behavioral perception of stop consonants are highly correlated, demonstrating the cABR’s behavioral relevance in speech perception. Finally, both behavioral perception and cABR are modulated by language background^46^.

**Figure 1.**
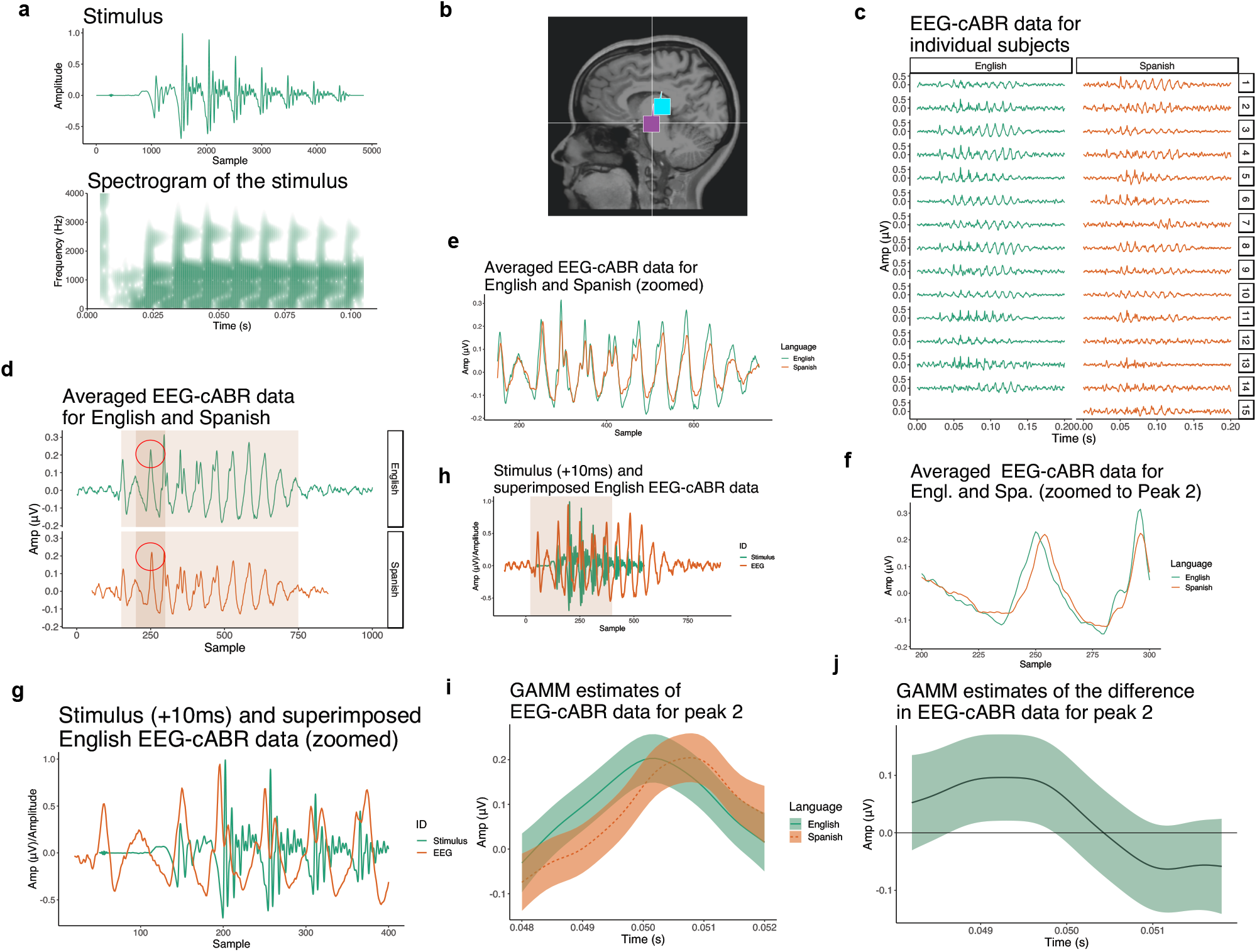
(**a**) Synthesized stimulus used in the cABR experiment with a spectrogram (0–4,000 Hz). This stimulus is also used in the computational experiments when the Generator is forced to output data with the objective to minimize the distance between generated data and the stimulus. The stimulus is played to subjects in the experiment. (**b**) The figure illustrates the dipole location of the onset peaks recorded during the cABR experiment for one speaker as localized in^46^ (magenta = peak 1, cyan = peak 2). The location suggests the recorded brain activity is indeed localized in the brain stem. Figure in (**c**) shows cABR recordings averaged for each subject across the 3,000 trials. (**d**) Individual subjects’ recordings are averaged for each language (with shades that indicate which parts are zoomed in in the following figures). Peak 2 is circled in red. (**e**) Zoomed cABR data for English and Spanish showing that most peaks (with the exception of peak 2) are almost perfectly aligned across the two languages. (**f**) Zoomed peak 2 showing peak latency differences between English and Spanish. Figure in (**h**) superficially parallels the stimulus with the cABR data. The brain signal in the experiment is manually delayed relative to the stimulus; for illustration, we manually aligned the two time-series by approximately aligning the burst of the stimulus and the first peak of the cABR data. Shaded part indicates the area zoomed in (**g**). (**i**) Predicted values of a Generalized Additive Mixed Model with Amplitude in *μ*V across time and the two languages (English vs. Spanish). For more details about the model, see Supplementary Table S2 and Section 2.3. (**j**) Difference smooth between English and Spanish cABR data. The area on the time scale (x-axis) in which the difference smooth’s confidence interval do not cross zero indicates significant difference in the cABR signal between the two languages.

The cABR data used in this paper comes from the previously published dataset in Zhao and Kulh^46^.^2^ The experiment measured the cABR when native English and Spanish subjects listened to a synthesized syllable, which was identified as /ba/ by English speakers and /pa/ by Spanish speakers. Data from a total of 15 Spanish and 14 English monolingual speakers were included in the analysis.

### 2.1 The stimulus

The stimulus is a CV syllable with a vowel /a/. The bilabial stop consonant has a Voice-Onset-Time (VOT) of +10ms and was synthesized by Klatt synthesizer in Praat software^64^. The syllable with 0ms VOT was first synthesized with a 2ms noise burst and vowel /a/. The fundamental frequency of the vowel /a/ began at 95Hz and ended at 90Hz. The silent gap (10ms) was then added after the initial noise burst to create syllables with the positive VOT. The waveform and spectrogram of the stimulus are shown in Figure 1a. The duration of the syllable is 100ms. Critically, monolingual English speakers identified the stimulus as /ba/ whereas native Spanish speakers identified the stimulus as /pa/, as reported in a previous behavioral experiment^46^. Individuals’ cABR were calculated by averaging across all available trials after standard preprocessing and trial rejection. Averaged values are visualized in Figure 1c. Further, the group-level cABR can be visualized by averaging over all subjects. The monolingual English group and the native Spanish group are represented in Figure 1d,e,f,g,h.

### 2.2 cABR data acquisition

The details of the recording methods can be found in Zhao and Kuhl^46^. Specifically, the cABR reported here is recorded using a traditional set-up of 3-EEG channels (i.e., CZ electrode on a 10-20 system, ground electrode on the forehead and the reference electrode on the right earlobe^45^). Two blocks of recordings (3,000 trials per block) were completed for each participants where trials were alternating in polarities.

### 2.3 A new statistical analysis

Zhao and Kuhl^46^ show that peak latency timing differs significantly for peak 2 between English and Spanish subjects using independent t-test. To analyze data with non-linear regression we fit the data averaged for each subject to Generalized Additive Mixed Models (GAMMs^65^) with the Amplitude of EEG-cABR in *μV* as the dependent variable and LANGUAGE (treatment-coded with English as level) as parametric term, a smooth for time, by-language difference smooth for time, and by-speaker random smooths as well as correction for autocorrelation (Figure 1i). The estimates of the model are in Supplementary Table S2. Even with random smooths included, the model features high degrees of autocorrelations. Significant difference does not arise for all windows of analysis likely due to correlation, but for a given window (from 240th to 260th sample), the difference smooth in Figure 1j suggest a significant difference in trajectory of the Amplitude between English and Spanish monolinguals in Peak 2 (*F* = 2.70, *p* = 0.015).

### 2.4 Results & interpretation of the cABR experiment

In summary, results from the cABR experiment demonstrated a robust effect of language background on the peak 2 latency of the cABR onset response. Particularly, the latency of peak 2, corresponding to the encoding of the onset of voicing, is significantly later in native Spanish speakers compared to the monolingual English speakers. Critically, the peak 2 latency was directly related to perception of the speech sound^46^. These suggest that the effect of language experience is reflected at very early stages of auditory processing, namely the auditory brainstem.

## 3 Computational Experiments

### 3.1 Model

We used the WaveGAN model^66^ (a DCGAN^11,67^ adaptation for audio) in our computational experiments. WaveGAN is a 1D deep convolutional generative adversarial model that operates directly on the waveform itself. The Generator *G* uses 1D transpose convolutions to upsample from the latent space *z*, while the Discriminator *D* uses traditional 1D convolutions to compute scores that assist in predicting the Wasserstein distance between the training distribution *x* and the distribution of generated outputs *G*(*z*)^68^. The architecture is outlined in Figure 2.

**Figure 2.**
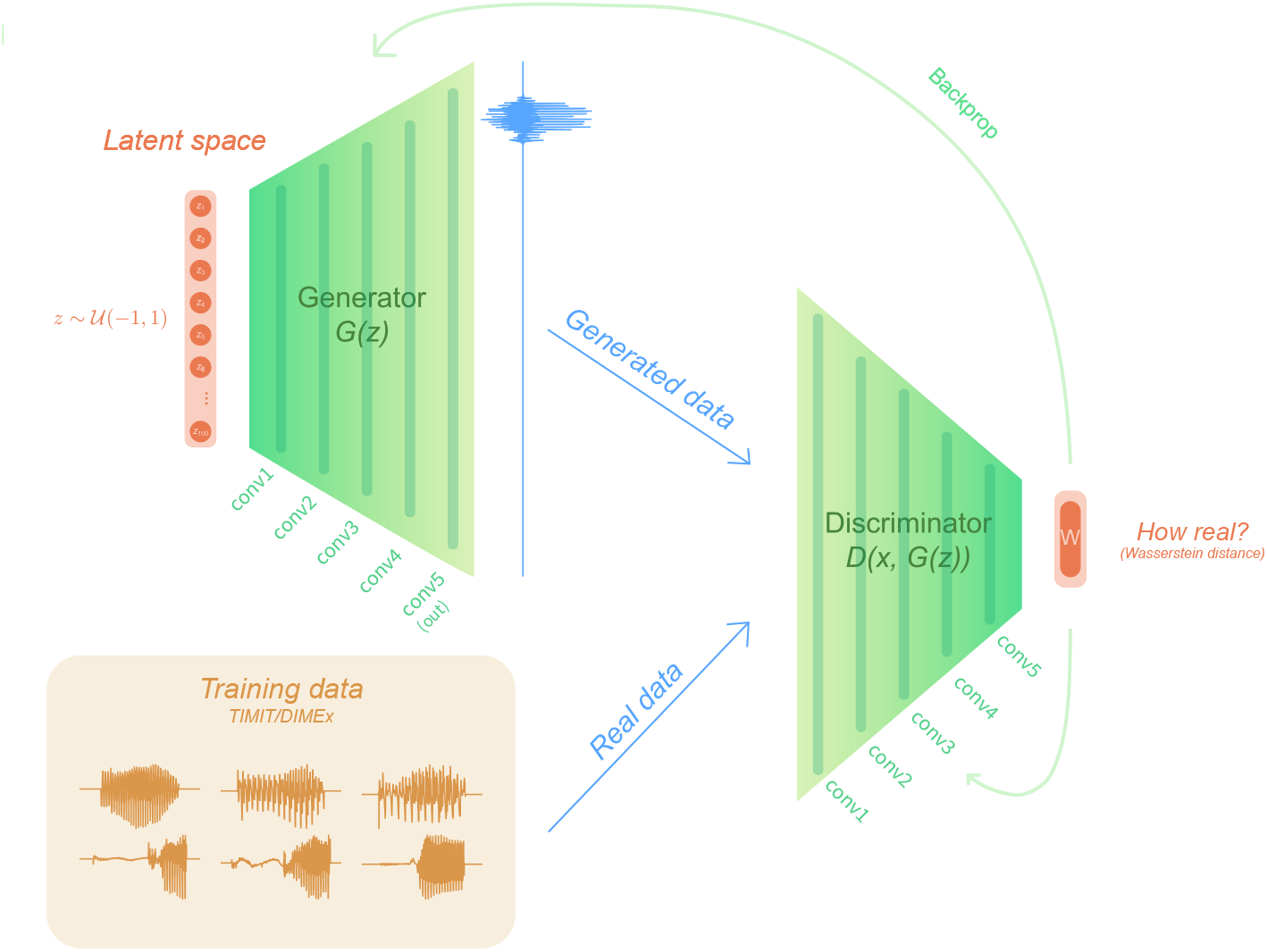
WaveGAN architecture^69^ (based on DCGAN^67^) used in training. The training data was taken from TIMIT and DIMEx as described in Section 3.2.

WaveGAN itself does not contain any visualization techniques. For analyzing and visualization of intermediate convolutional layers, we use a visualization technique for the Generator’s^50^ and the classifier’s internal layers^51^ (which has almost identical structure to the Discriminator). In Beguš and Zhou^50,51^, we argue that averaging over feature maps after ReLU activation yields a highly interpretable time series data for each convolutional layer that summarizes what acoustic properties are encoded at which layer.

For these experiments, we set *z* to be a 100-dimensional vector (following the WaveGAN proposal^66^), which the Generator projects into a 2D tensor that is passed through 5 transpose convolutional layers, ending in an audio output of 16,384 samples. The Discriminator similarly is composed of 5 (traditional) convolutional layers, with a hidden layer at the end that outputs the Wasserstein metric. No optimization was done over the number of convolutional layers nor any other part of the model or training configuration; we took the default 5-layer configuration of WaveGAN/DCGAN with a 16,384 sample output^66,67^. The choice of the number of convolutional layers does not substantially alter the encoding of acoustic features across layers^50^. The Discriminator also makes use of a process called phase shuffle^66^, which applies random perturbations to the phase of each layers’ activations to prevent the Discriminator from accessing periodic artifacts characteristic of transpose convolutions.

### 3.2 Data

Spanish training data was taken from the DIMEx100 corpus^49^. This dataset consists of audio recordings of 5010 sentences in Mexican Spanish, recorded from 100 speakers mostly from around Mexico City. The dataset is balanced in gender and represents primarily the Mexico City variety of Spanish^49^. English training data was taken from the TIMIT speech corpus^48^. The TIMIT dataset contains recordings of 6300 sentences of American English, spread across 8 dialects and 630 speakers^48^.

For the purposes of training, we slice the first syllable from words that begin with a voiced or voiceless stop.^3^ Specifically, we slice sequences of the form #CV, where # represents a word boundary, C represents a voiced or voiceless stop, and V represents a vowel. For both English and Spanish, the voiceless stops consist of [p, t, k] and the voiced stops consist of [b, d, g]. The number of sequences beginning with each stop in both datasets are shown in Table 1. The relative frequencies of phonemes differ across TIMIT and DIMEx datasets, but this is desirable even if some sequences are overrepresented as we aim to train our network on data that most closely resembles actual distributions of phonemes in English and Spanish.

**Table 1.** Counts of sequences beginning with each stop for each corpus.

|  | p | t | k | b | d | g |
| --- | --- | --- | --- | --- | --- | --- |
| TIMIT | 1018 | 1799 | 2112 | 1789 | 2530 | 673 |
| DIMEx100 | 3015 | 1808 | 5558 | 1477 | 8023 | 478 |

### 3.3 Training

We trained the DIMEx100 model for approximately 38,649 steps, after which mode collapse was observed. To match the two models in the number of steps, we train the TIMIT model for 40,730 steps. To replicate the results and to control for idiosyncracies of individual models, we trained one additional TIMIT and one additional DIMEx100 model (for 41,818 and 39,417 steps, respectively).

### 3.4 Generating outputs that approximate the stimulus

In order to test the stimuli against the Generator network, we use latent vector recovery techniques^41,42^ to find the latent variables that result in outputs closest to the stimuli. We then generate outputs using these latent variables and analyze each layer of the network given that latent space. This is a novel approach to paralleling representations in deep neural decoder networks and brain imaging outputs: the model’s internal representations are chosen such that the generated output maximally resembles the stimulus in the brain experiment. Norman-Haignere and McDermott^43^ propose a somewhat similar procedure, where outputs of the brain experiments are paralleled with synthetic stimuli “designed to yield the same responses as the natural stimulus”^43^. In our case, the directionality of forced input is reversed: we seek internal representations that result in maximal matching between the actual stimulus and the model’s output.

We use gradient descent with stochastic clipping^41^, on the mean absolute error of the spectrogram of the stimulus and the spectrogram of the generated output^42^. We sample many random latent vectors uniformly for consistency, and optimize using the ADAM optimizer with learning rate of 1e *−* 2, first moment decay of 0.9, and second moment decay of 0.99. We optimize for 10,000 steps, after which the majority of outputs converge. We adapt the objective function from Keyes et al.^42^ (listed below, where *G* is the generator network, *S* takes an audio signal to a spectrogram, and *s* is the target stimulus):

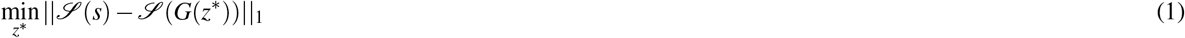

As the Generator generates a fixed-length output, we must zero-pad the target stimulus before performing loss computations. Interestingly, while all training samples were simply right-padded to the desired dimension, we found that introducing varying amounts of left-padding had differing results in the quality of the generation. The DIMEx100 model, in particular, is extremely sensitive to the left pad, creating nonsense forced outputs with a left pad of 0 samples and creating much closer outputs with a left pad of 1000 samples. The TIMIT model is much less sensitive to the pad, and generates fairly close samples with a left pad of anything from 0 to 1000 samples. This difference may be due to differences in the slice distribution of the two corpora, but for the sake of consistency we used a left pad of 1000 samples for both models.

### 3.5 Procedure

The visualization technique in Beguš and Zhou^50,51^ allows us to test acoustic representations of intermediate convolutional layers of both the Generator that mimics the production principle in speech and the Discriminator that mimics the perception principle. Here, we compare the outputs of the proposed technique to outputs of brain imaging experiments.

The relationship between the stimulus played to the subjects in the cABR experiment and the amplitude of the cABR recording is paralleled to the relationship between the generated outputs forced to resemble the stimulus and the fourth (second to last) convolutional layer in the Generator network.

To extract interpretable data from intermediate convolutional layers in the computational experiment, we force the Generator to output #CV syllables that most closely resemble the stimulus used in the cABR experiment (Figure 1a), as described in Section 3.4.

The Generator mimics the production aspect of speech. The cABR experiment, however tests encoding of a phonological contrast in the perception task. For this reason, we also test the relationship between the input to the Discriminator (that mimics speech perception) and its corresponding first convolutional layer.

The inputs to the Discriminator are two fold: first we feed the Discriminator the raw waveform of the actual stimulus used during the brain experiment (Figure 1a). This means that the CNN and the brain during the actual experiment are tested on the exact same data. This experiment reveals a high degree of similarities between the signals even when no transformations are performed. However, this experiment only yields one observation per trained model (four total) which prevents an inferential statistical analysis. To test learned representations in the Discriminator further and to increase variability of its representation, we also feed it the Generator’s outputs forced to resemble the stimulus (according to Section 3.4).

The fourth and first convolutional layers, respectively, are analyzed by averaging over all feature maps after ReLU or Leaky ReLU activations^50,51^. This results in a time series *t* for each Convolutional layer as in (2) from Beguš and Zhou^51^.^4^

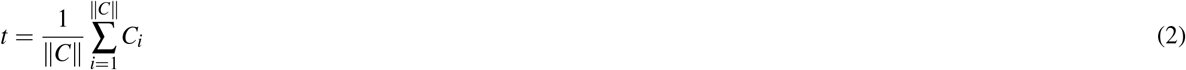

The cABR experiment suggest that peak 2 latency differs significantly between English and Spanish speakers^46^. To test encoding of the same acoustic property in intermediate convolutional layers, we measure peak latency timing between amplitude peaks in output/input and amplitude peaks in the second to last or first convolutional layer (Conv4 or Conv1) in the Generator and Discriminator networks, respectively.

To extract peak timing in each layer, we first generate 20 Generator outputs per model that are forced to resemble the stimulus (according to Section 3.4). In three outputs of the first and the second TIMIT replication we were unable to identify the periodic structure, which is why they were removed from the analysis. A total of 74 forced outputs were thus created (2 replications of TIMIT and DIMEx trained models each). For each generated output, we obtain the corresponding representations in the fourth (immediately following) convolutional layer (Conv4) as described in Section 3.4 by averaging over all feature maps. This yields a time-series data. The generated outputs and the time-series data from the fourth convolutional layers (upsampled) are then annotated for vocalic periods. Peak timing for each vocalic period is obtained in Praat^64^ with parabolic interpolation. Because the values of the fourth convolutional layer can only be positive due to the ReLU transformation, we also take absolute values of the waveforms for the comparison. It appears that the preceding convolutional layers closely follow amplitude changes in the output layer. Converting waveforms into absolute values is necessary in order to capture peak activity of negative values as well as positive values of waveforms (ReLU can only be positive). This step also reduces effects of different recording condition of the two corpora (TIMIT and DIMEx).

Peak latency (Δ*t*_*n*_) was calculated as a difference in timing between the peak of absolute values of the output 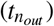 and the peak of the fourth convolutional layer 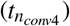 in the Generator.

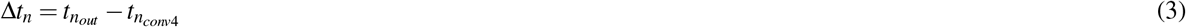

The burst is annotated as the 0th period and every consecutive period as *n*th period. The burst is not saliently present in all outputs. A total of 51 bursts (=0th period) were included in the analysis.

To test peak latency in the Discriminator network, we feed the Discriminator the actual stimulus as well as the same 74 generated outputs from the Generator forced to resemble the stimulus. Peak latency in the Discriminator was calculated as a difference in timing between the absolute value of peak of the input and the peak of the first (immediately following) convolutional layer. The same annotations as for output-Conv4 analysis in the Generator were used to extract peak timing from the forced generated outputs and the first convolutional layer (Conv1) in the Discriminator network (according to (3)).

## 4 Results

### 4.1 Similarities in encoding

First, we observe that the cABR signal and the output of the fourth/first convolutional layers are highly similar. The computational experiment that most closely resembles the cABR experiment is when the Discriminator gets the actual stimulus as the input. Figure 3 parallels the cABR response averaged across subjects and the response in the first convolutional layer of the Discriminator averaged across replications. The two modalities show almost exactly the same response to the stimulus with highly similar shapes of periods.

**Figure 3.**
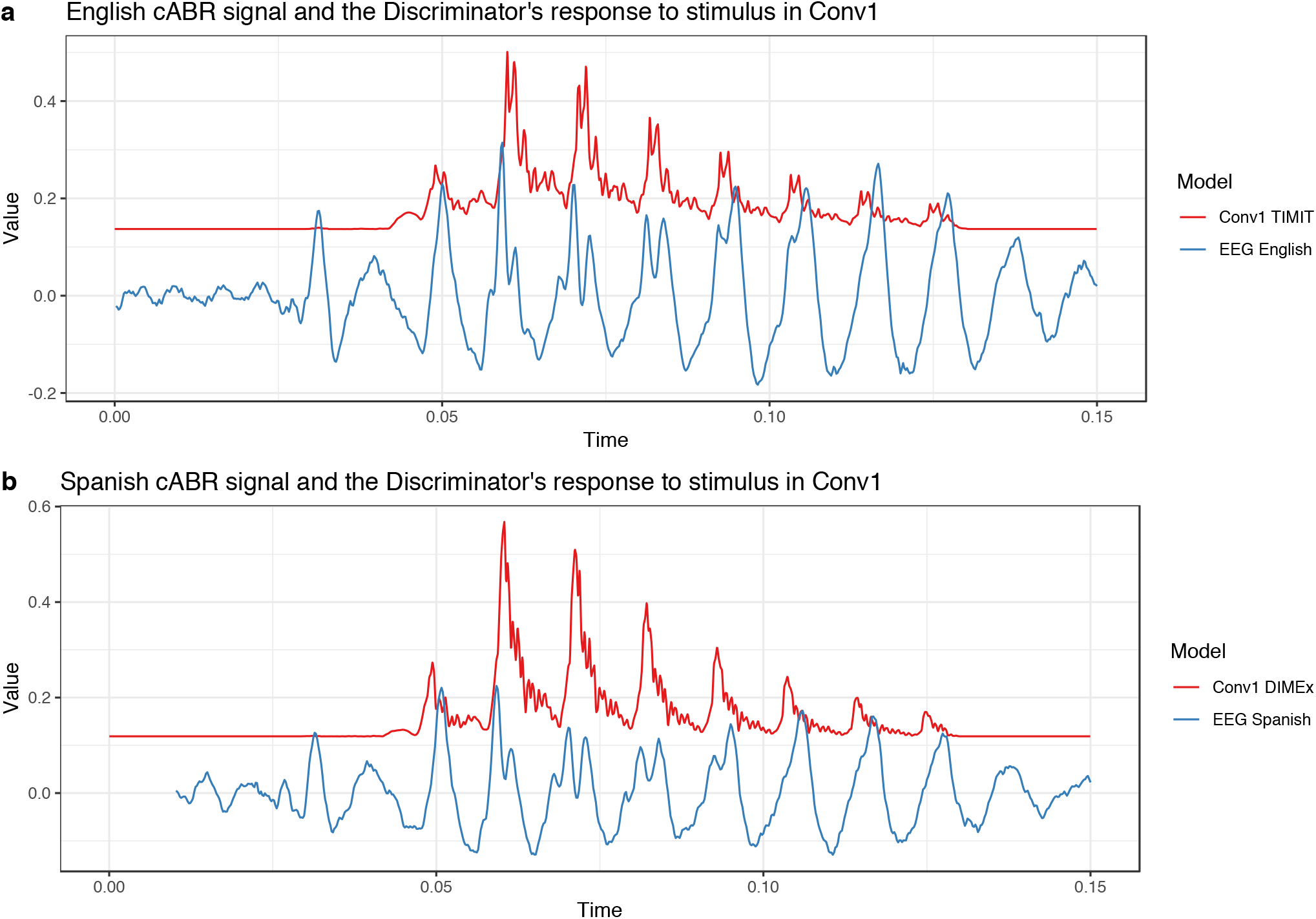
(**a**) Values of the English cABR experiment averaged across subjects (blue) and values of the first convolutional layer in the TIMIT-trained model (Conv1 TIMIT) averaged across replications when the Discriminator gets the actual stimulus as the input (red). The two time series are aligned such that the peaks corresponding to the burst have the same timing. The values of the Conv1 signal are increased 50-times for comparison. (**b**) Values of the Spanish cABR experiment averaged across subjects (blue) and values of the first convolutional layer in the DIMEx-trained model (Conv1 DIMEx) averaged across replications when the Discriminator gets the actual stimulus as the input (red). The two time series are aligned such that the peaks corresponding to the burst have the same timing. The values of the Conv1 signal are increased 50-times for comparison.

To quantify this observation, we perform dynamic time warping (DTW) on the two time series aligned at the peak of the burst and compute Pearson’s product-moment correlation (*r*) between the two time-series when the convolutional signal is increased 33.3-times or 50-times, decreased such that silence reaches 0, and downsampled with linear interpolation (to match the sampling of the cABR signal). For the period between the peak of English burst to the 130th milisecond, the correlation coefficient for English is *r* = 0.74 and Spanish *r* = 0.77. At the individual period level, the correlation is even higher. For example, between the 80th and 90th miliseconds which captures 2 periods, the correlation coefficient for English is *r* = 0.90 and Spanish *r* = 0.82.

The technique to parallel biological and artificial neural responses to spoken language inputs allow us to go beyond comparing the two signals in terms of correlations and allows us to analyze encoding of actual acoustic properties. We focus on peak latency because it has been shown in cABR experiments that English and Spanish monolinguals differ significantly in this property^46^.

Visualizations of raw peak latency timing in the Generator and the two Discriminator experiments (in Figures 3, 4g,h, and 6g,h) suggest that there is a consistent timing difference between the TIMIT-trained models (English) and the DIMEx-trained models (Spanish). The peak latency (Δ*t*_*n*_) is more negative in the Spanish-trained models compared the English-trained models in a few periods. In other words, the peak activity in the Spanish-trained models occurs later compared to the English-trained models, which is consistent with the results of the brain experiment. This observation is consistent in both the Generator and the Discriminator as well as across replications.

**Figure 4.**
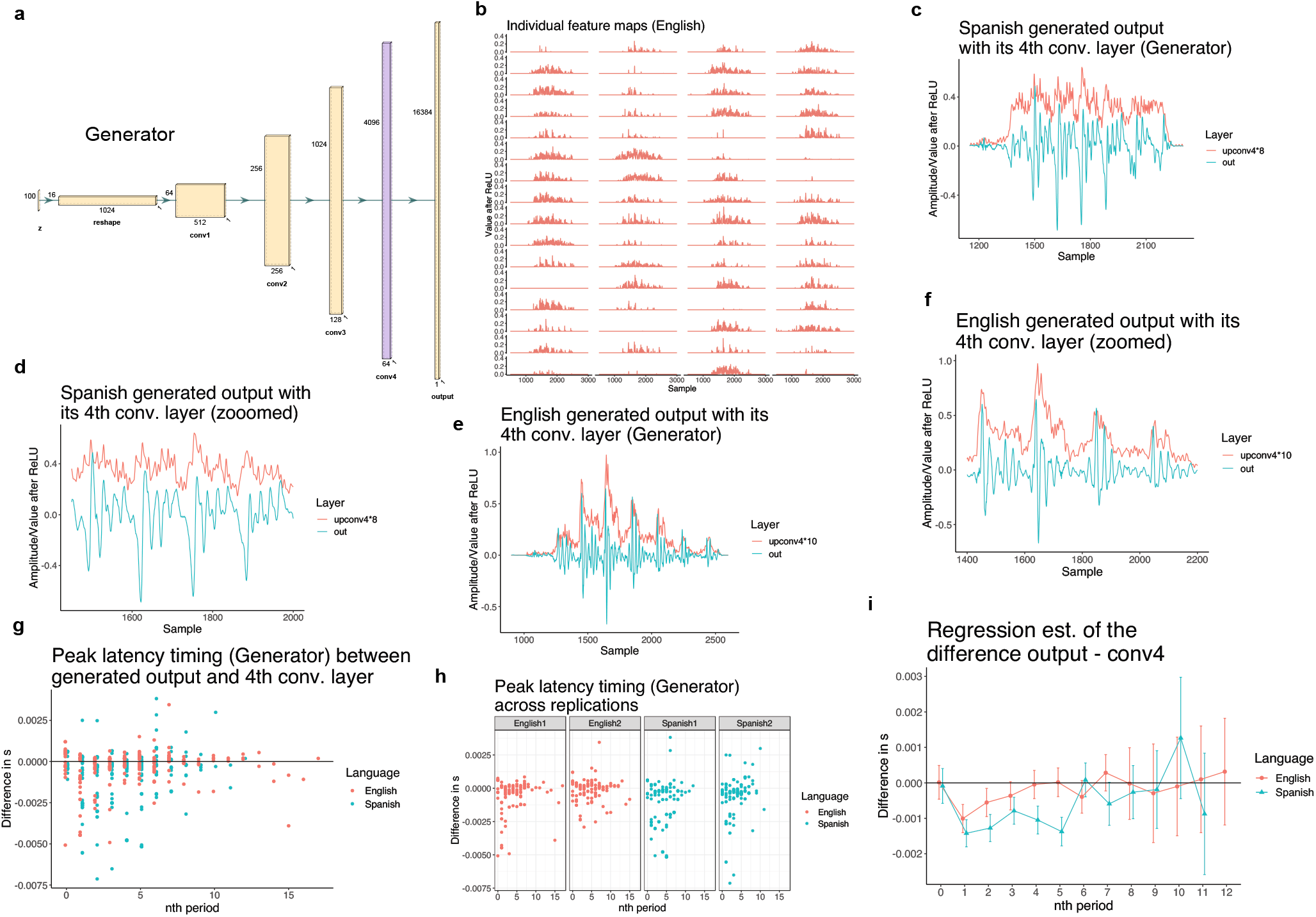
(**a**) The structure of the Generator network with five convolutional layers^66^. The fourth convolutional layer (Conv4; second to last) is color-coded with purple. (**b**) All 64 individual feature maps for a single output forced to closely resemble the stimulus from the fourth convolutional layer (Conv4) after ReLU (upsampled). (**c**) One Spanish output (in green) forced to resemble the stimulus with the corresponding values from the fourth convolutional layer (Conv4) averaged over all feature maps. The plot illustrates peak latency between output and Conv4 for the burst and each vocalic period. (**d**) A zoomed version of (d) focusing on four vocalic periods. (**e**) One English output (in green) forced to resemble the stimulus with the corresponding values from the fourth convolutional layer (Conv4) averaged over all feature maps. The plot illustrates peak latency between output and Conv4 for the burst and each vocalic period. (**f**) A zoomed version of (f) focusing on four vocalic periods. (**g**) Raw peak latency timing (output peak time - Conv4 peak time) for burst (=0) and each nth vocalic period across the two conditions (English vs. Spanish). Periods above the 12th period are rare and are discarded from the statistical analysis due to a small number of attestations. The data is pooled across the two replications. (**h**) Raw peak latency timing across the replications (first and second replication) and two conditions (English and Spanish). (**i**) Linear regression estimates for the peak latency timing between the two conditions (English vs. Spanish). Periods above the 12th period are discarded from the analysis due to a small number of attestations. The data is pooled across the two replications.

### 4.2 Peak latency: the Generator

To test the significance of the peak latency differences in the Generator, we fit the data from the 74 forced outputs (Section 3.5) to a linear regression model with the PEAK LATENCY timing as the dependent variable and three predictors: LANGUAGE, nTH PERIOD, and REPLICATION with all two-way and three-way interactions.

The LANGUAGE predictor has two levels (English and Spanish) and is treatment-coded with English as the reference level. The nTH PERIOD predictor has 13 levels (for each period and the burst) and is treatment-coded with 1st period as the reference level. Periods above the 12th period are discarded from the analysis due to a small number of attestations (see Figures 4 and 6). REPLICATION is sum-coded with two levels (first and second).

Estimates of the model are given in Supplementary Table S3 and Figure 4i. Pairwise comparisons in Table 2 reveal that peak timing does not differ significantly for the burst (0th period) and the first period, but the difference becomes significant for 2nd, 4th, 5th, and 7th periods (see all estimates in Table 2). If we adjust pairwise comparisons with False Discovery Rate (FDR) adjustment, only differences for the 4th, and 5th period are significant (*p*-value for the 2nd period is 0.0501). A subset of peak latency differences are significant in individual replications too (See Supplementary Figure S5). For example, in the second replication, peak latency is significantly different in the 1st period (*β* = 0.0012, *d f* = 517, *t* = 2.890, *p* = 0.03 with FDR adjustment).

**Table 2.**
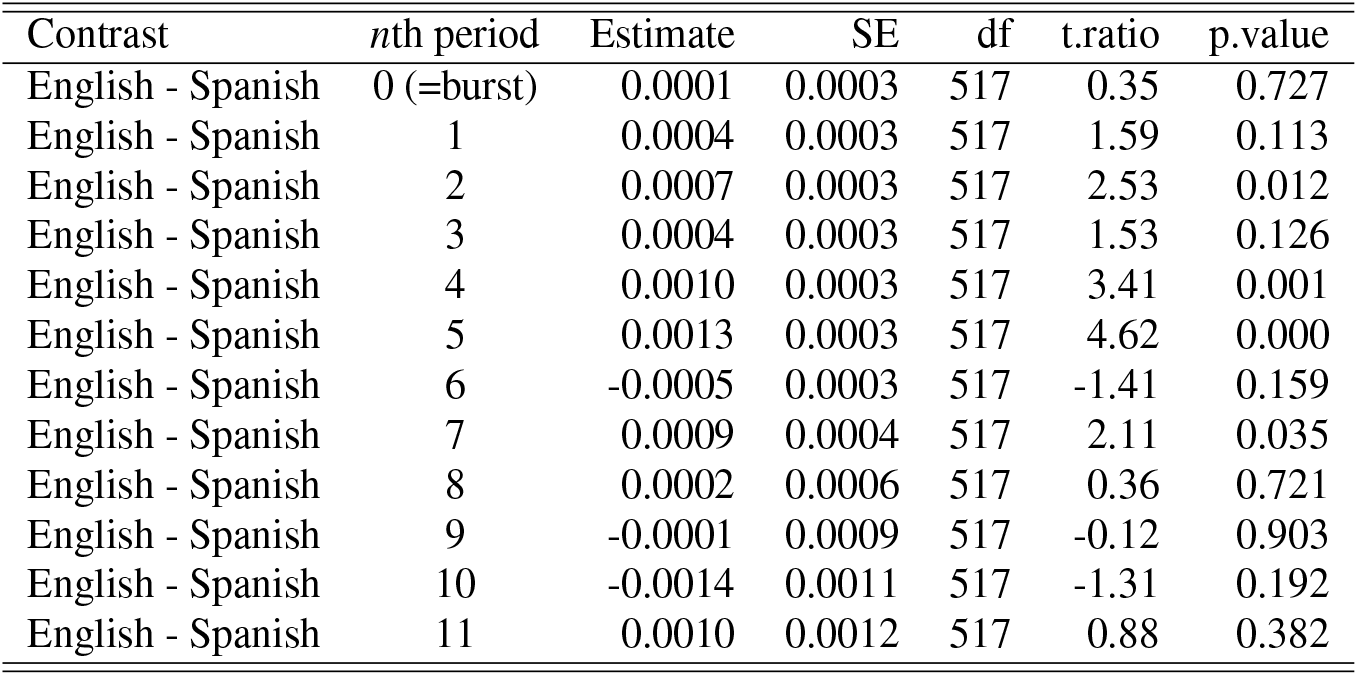
Pairwise contrasts in peak timing difference between English and Spanish (despite significant interactions pooled across replications) in the Generator network (with *emmeans* package^72^). The burst is marked by the 0th period. The 12th period is not estimated due to lack of data.

### 4.3 Peak latency: the Discriminator with the stimulus

To analyze peak latency in the Discriminator network, we first feed the Discriminator network the raw waveform of the stimulus used in the brain experiment. Using the averaging technique (in 2) on the first convolutional layer (Conv1), we get one corresponding time series data per each model (four total: TIMIT1, TIMIT2, DIMEx1, DIMEx2). We can parallel these time-series data and analyze the timing of peak activity per period in each model. In Figure 5, we averaged values across replications in a similar way as we averaged values over the subjects in the cABR experiment in Figure 1. The results suggests that in the first period, the peak activity in the English-trained model precedes the Spanish-trained model, parallel to the brain activity in peak 1 of the cABR experiment. Averaged peak timing in the English-trained model precedes the Spanish-trained model also in peaks 2, 4, 6, and 7. Even the magnitude of this effect are similar across the cABR and convolutional signals: in cABR, the English peak precedes the Spanish peak by 0.9 ms; in the convolutional layers of the Discrimninator, the peak in the TIMIT-trained model precedes the peak in the DIMEx-trained model by 0.5 ms.

**Figure 5.**
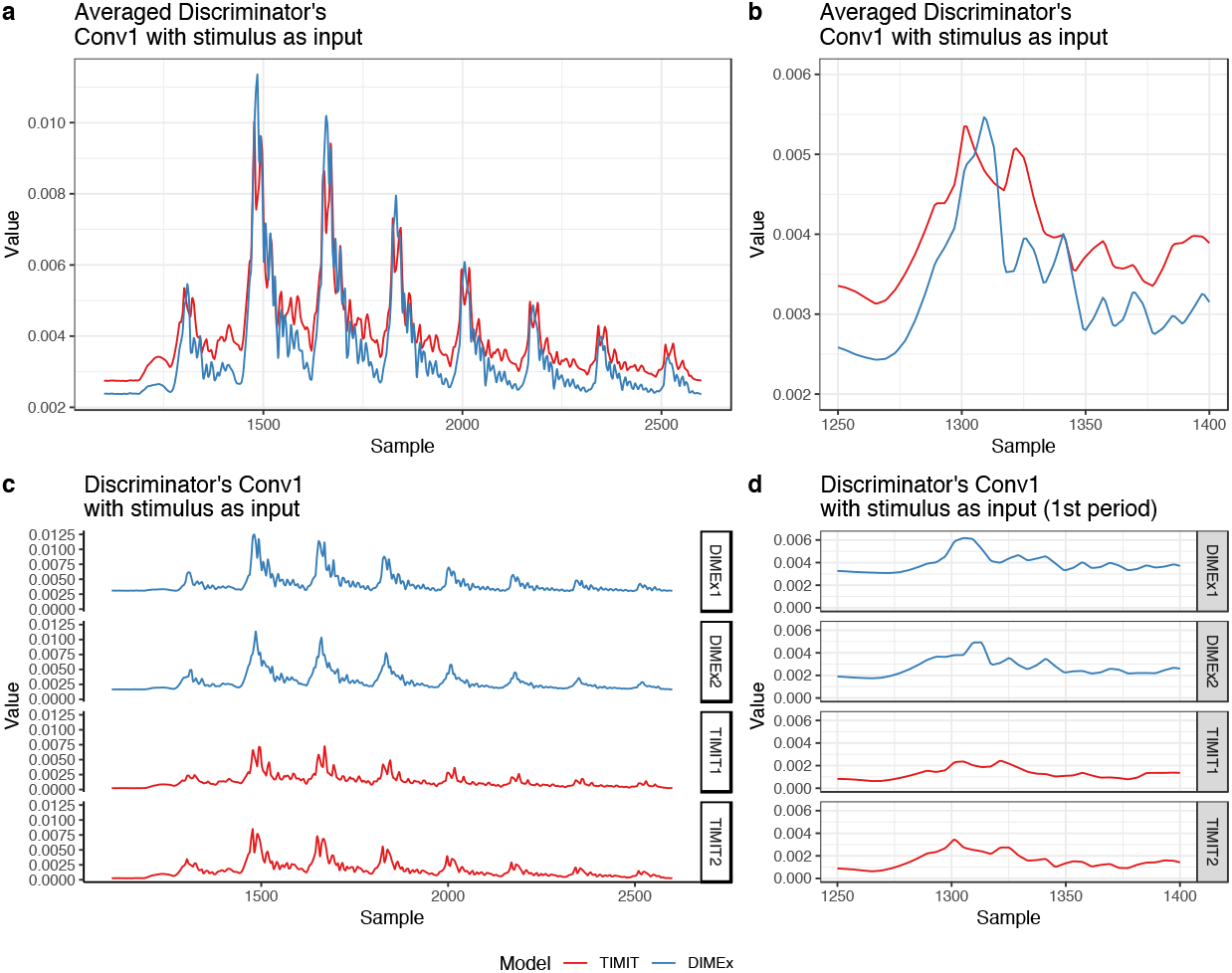
(**a**) Values of the first convolutional layer (Conv1) when the stimulus used in the brain experiment is the input to the Discriminator network. The figure shows averaged values across two replication of each models (DIMEx-trained or Spanish and TIMIT-trained or English). Values of the TIMIT outputs were increased by +0.0025 on the y-axis to facilitate the comparison between signals. (**b**) Averaged values of the first convolutional layer (Conv1) when the stimulus used in the brain experiment is the input to the Discriminator network, zoomed in to the first period. Values of the TIMIT outputs were increased by +0.0025 on the y-axis to facilitate the comparison between signals. (**c**) Individual values for each model (TIMIT1, TIMIT2, DIMEx1, DIMEx2) of the first convolutional layer (Conv1) when the stimulus used in the brain experiment is the input to the Discriminator network. (**d**) Individual values for each model (TIMIT1, TIMIT2, DIMEx1, DIMEx2) of the first convolutional layer (Conv1) when the stimulus used in the brain experiment is the input to the Discriminator network, zoomed in to the first period.

Paralleling the Discriminator’s responses to the actual stimulus approximates the brain experiment most closely. This approach, however, does not allow for inferential statistical tests on the distributions due to the small number of obtained samples. In order to analyze learned representations of the Discriminator’s internal convolutional layers and perform inferential statistical tests on the distributions, we also feed the Discriminator the outputs from the Generator forced to resemble the stimulus (according to the procedure described in Section 3.4).

### 4.4 Peak latency: the Discriminator with generated outputs

To test the significance of peak latency differences in the Discriminator network when it is fed the Generator’s forced outputs, we perform the same statistical procedure as described in Section 4.2. The peak latency (Δ*t*_*n*_) for the *n*th period in the Discriminator is calculated as the difference between the absolute peak timing of the input and peak timing of the first convolutional layer (Conv1) for each period. The model in Figure 6i and Supplementary Table S4 includes LANGUAGE, nTH PERIOD, and REPLICATION (coded as in Section 4.2) and all interactions as predictors and the peak latency timing as the dependent variable. The pairwise comparisons are in Table 3. Peak latency for the burst (=0th period) does not differ significantly across the two languages. The difference is significant for the 3rd and the 6th period. (also when tested with FDR correction). The 6th period in the Discriminator is the only period in which peak latency timing is significant in the opposite direction than all the other trends in the Generator and the Discriminator. A subset of peak latency differences are significant in individual replications too (See Supplementary Figure S6).

**Figure 6.**
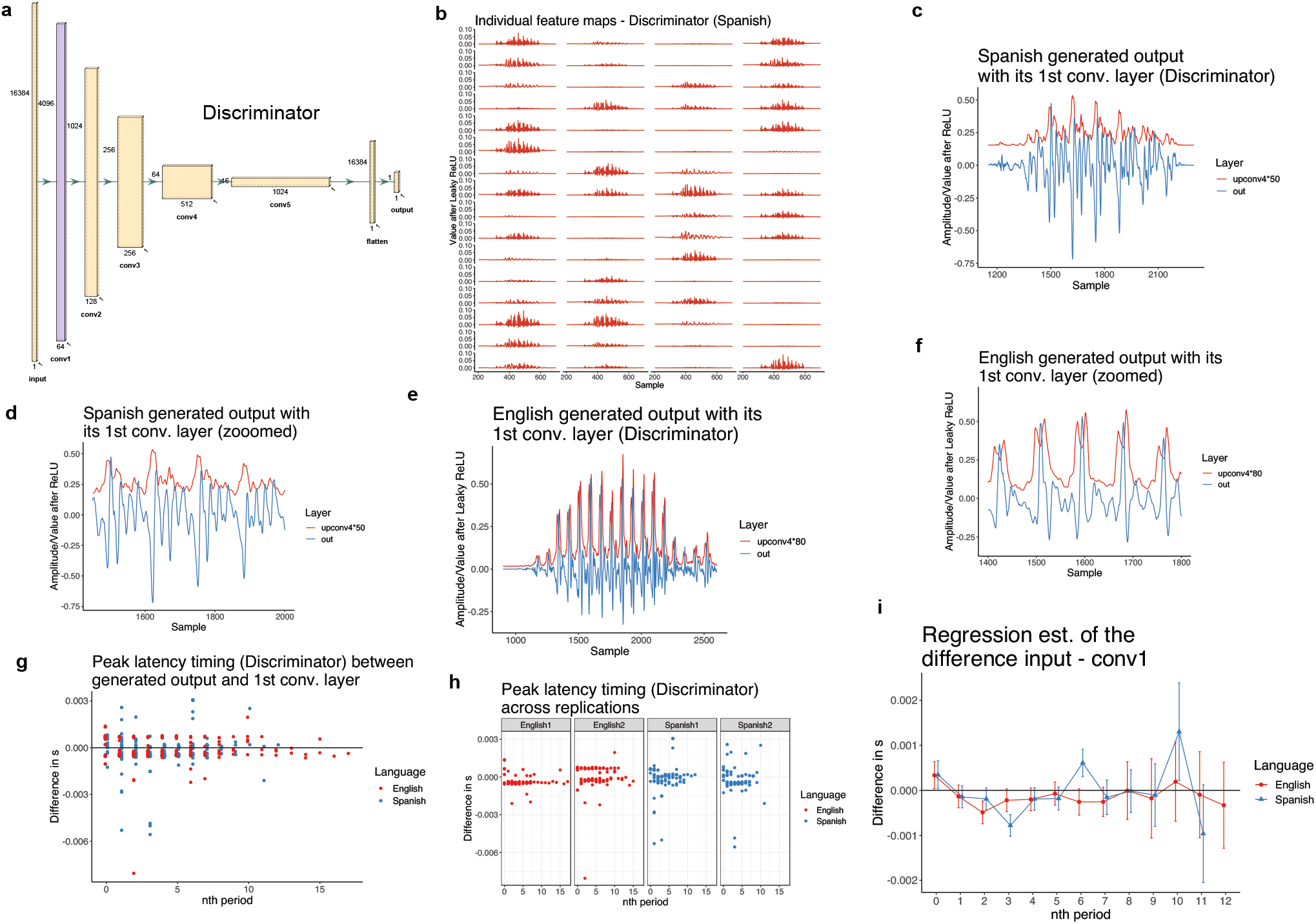
(**a**) The structure of the Discriminator network with five convolutional layers^66^. The first convolutional layer (Conv1; second to last) is color-coded with purple. (**b**) All 64 individual feature maps for a single input (the Generator’s forced output) from the first convolutional layer (Conv1) after Leaky ReLU. (**c**) One Spanish input (in blue) from the Generator’s forced output with the corresponding values from the first convolutional layer (Conv1) averaged over all feature maps. The plot illustrates peak latency between input and Conv1 for the burst and each vocalic period. (**d**) A zoomed version of (d) focusing on four vocalic periods. (**e**) One English input (in blue) from the Generator’s forced output with the corresponding values from the first convolutional layer (Conv1) averaged over all feature maps. The plot illustrates peak latency between input and Conv1 for the burst and each vocalic period. (**f**) A zoomed version of (e) focusing on five vocalic periods. (**g**) Raw peak latency timing (input peak time - Conv1 peak time) for burst (=0) and each nth vocalic period across the two conditions (English vs. Spanish). Periods above the 12th period are rare and are discarded from the statistical analysis due to a small number of attestations. The data is pooled across the two replications. (**h**) Raw peak latency timing across the replications (first and second replication) and two conditions (English and Spanish). (**i**) Linear regression estimates for the peak latency timing between the two conditions (English vs. Spanish). Periods above the 12th period are discarded from the analysis due to a small number of attestations. The data is pooled across the two replications.

**Table 3.** Pairwise contrasts in peak timing difference between English and Spanish (despite significant interactions pooled across replications) in the Discriminator network (with *emmeans* package^72^). The burst is marked by the 0th period. The 12th period is not estimated due to lack of data.

| Contrast | nth period | Estimate | SE | df | t.ratio | p.value |
| --- | --- | --- | --- | --- | --- | --- |
| English - Spanish | 0 (=burst) | -0.0000 | 0.0002 | 517 | -0.00 | 0.999 |
| English - Spanish | 1 | 0.0000 | 0.0002 | 517 | 0.05 | 0.960 |
| English - Spanish | 2 | -0.0003 | 0.0002 | 517 | -1.63 | 0.103 |
| English - Spanish | 3 | 0.0006 | 0.0002 | 517 | 3.24 | 0.001 |
| English - Spanish | 4 | 0.0000 | 0.0002 | 517 | 0.10 | 0.923 |
| English - Spanish | 5 | 0.0001 | 0.0002 | 517 | 0.74 | 0.462 |
| English - Spanish | 6 | -0.0008 | 0.0002 | 517 | -3.81 | 0.000 |
| English - Spanish | 7 | -0.0001 | 0.0003 | 517 | -0.31 | 0.756 |
| English - Spanish | 8 | 0.0000 | 0.0004 | 517 | 0.03 | 0.977 |
| English - Spanish | 9 | -0.0001 | 0.0006 | 517 | -0.11 | 0.913 |
| English - Spanish | 10 | -0.0011 | 0.0007 | 517 | -1.61 | 0.108 |
| English - Spanish | 11 | 0.0009 | 0.0007 | 517 | 1.28 | 0.203 |

The magnitude of the effect of language is similar across the cABR and convolutional layer signals. In the cABR signal, the peak 2 timing difference between English and Spanish monolinguals is 0.9 ms. In the experiments with generated data on the Generator and the Discriminator (Sections 4.2 and 4.4), the peak latency (output - Conv1) differs between TIMIT-trained and DIMEx-trained models in the range from 0.6 to 1.3 ms (for those results that are significant).

Peak latency differences are consistently significant between Spanish-trained and English-trained models even if waveforms are not converted into absolute values (Supplementary Materials Section 2). In such case, however, the Spanish-trained models show more positive peak latency timing. Nevertheless, the fact that significant results persist even in tests that do not include absolute values suggest that there are robust differences in peak latency timing in the second-to-last convolutional layers between Spanish and English-trained models that persist even with different analytical choices.

## 5 Discussion

The paper presents an interpretable technique that allows paralleling biological and artificial neural representations and computations in spoken language. The results of the proposed technique suggest that the speech input is represented in a highly similar way in the two signals and that peak latency in intermediate activations in English-trained and Spanish-trained deep convolutional networks differ in similar ways as the peak latency in cABR signal of English and Spanish speakers.

### 5.1 Similarities

Paralleling the response of intermediate convolutional layers with the brain stem’s response to the exact same stimulus reveals a high degree of similarities between the two signals. The shapes of periods responding to the two signals are almost identical and they also match in timing. These similarities arise without any transformations between the signals. To our knowledge, the cABR and convolutional layer response to the same syllable are the most similar brain and ANN signals reported thus far that require no linear transformations.

To move beyond comparing similarities, we also analyze differences in encoding of specific phonetic properties between the two signals. Peak latency has long been a focus of cABR studies^46,73,74^. Zhao and Kuhl^46^ argue that peak 2 latency differs significantly based on language experience, where the two different languages (Spanish and English) have substantially different encoding of a phonetic property which is correlated to the perception of the sound across individuals: voiceless (e.g. [ta]) vs. voiced (e.g. [da]) sounds. This suggests that phonetic features that represent a phonological contrast in language can be encoded early in the auditory pathway—already in the brainstem.

Peak latency is an interpretable feature that can be analyzed with standard acoustic methods in deep convolutional networks. We analyze peak latency encoding with a technique for visualization of intermediate convolutional layers^50,51^ that uses summation to identify peak activity in intermediate convolutional layers relative to the input/output. Because encoding of VOT duration in the form of peak latency appears to be present already at the brain stem level (based on the cABR experiment), we conduct the comparison of peak latency on the immediately preceding — second-to-final — convolutional layer relative to the input/output and parallel this information to the cABR signal in the brain stem relative to the stimulus.

The results of the computational experiment suggest that peak amplitude timing of the second to last convolutional layer relative to the speech input/output do not differ significantly for the burst, but do differ significantly for consecutive vocalic periods based on the language of training data: English (with long VOT encoding of voicing in stops) and Spanish (without long VOT encoding of voicing stops). The difference in timing of peaks (peak timing in audio input/output minus peak timing of second-to-last convolutional layer per each vocalic period) is significantly more negative in Spanish-trained model compared to the English trained model for several periods following the burst. The difference is significant both in the Generator (the production principle) as well as in the Discriminator (the perception principle). The peak latency also operates in the same direction across the two replications (in eight models total) with only one exception, which suggests the results are not an idiosyncratic property of individual models.

The results suggest that a highly interpretable acoustic property — peak latency — that indicates peak activity in the brain stem and in the intermediate convolutional layer relative to the stimulus/input/output based on a common operation, summation/averaging of the signal, is encoded in similar ways both in the earlier intermediate convolutional layers and the cABR signal. The encoding is similar both in the direction of latency (English peaks precede Spanish peaks) as well as in magnitude (0.9 ms in cABR vs. 0.5–1.3 ms in convolutional layers).

The only notable difference between the cABR experiment and the convolutional layers is that only the first period differs significantly in the cABR experiment, while multiple periods have significant peak timing differences in the convolutional layers (beginning with the second period in the Generator). It is possible that the subsequent periods in convolutional layers show significant peak timing differences because their signal are substantially stronger (higher amplitudes) compared to the first period. Noise in the output can have a more substantial effect on the results more when signal-to-noise ratio is low, i.e. in the first period in the convolutional layers.

A more conservative conclusion based on the results is that encoding of speech signal, and more specifically, of peak timing, can in general differ according to the language exposure (English vs. Spanish) in similar ways between the intermediate convolutional layers and the brain stem. Under this interpretation, it is possible that the similarities observed between cABR and convolutional layers have different underlying causes. For example, it is possible that peak latency in cABR is caused by VOT differences, while peak latency in the convolutional layers are caused by general vocalic encoding that differ across languages. Even in such a case, the conclusion that the cABR and convolutional layers response to the speech signal in highly similar ways remains.

Under a less conservative reading of the results, the difference in VOT encoding causes the peak latency differences both in the brain stem and in convolutional layers. Peak latency for burst is not significant neither in the brain nor in the intermediate convolutional layers, while subsequent periods show a significant difference in timing in both modalities. The magnitude of the timing as well as the direction of differences are the same across both modalities.

### 5.2 Causes of similarities

The results in this paper raise a question of what properties of deep convolutional networks and the cABR signal cause the similarities in encoding of an acoustic phonetic property. The main mechanism behind the technique for analyzing acoustic properties in intermediate convolutional layers is a simple averaging of activations across individual feature maps (in equation 2). The second to last convolutional layer (Conv4 in the Generator and Conv1 in the Discriminator) has 64 filters which result in 64 feature maps for each input/output. Individual feature maps offer limited interpretability, but a simple averaged sum over all feature maps after ReLU or Leaky ReLU activation offers highly interpretable time series data^50,51^.

Similar to this proposed computational technique, cABR data represents a summation of neural activity in the brain stem (and potentially also from other non-subcortical sources)^75,76^. The basic principle for obtaining the signal in both the brainstem and intermediate convolutional layers is thus similar: averaging of individual neural activity (biological and artificial) across the time domain.

Based on these similarities, it is reasonable to assume that both signals represent at least superficially similar computations. Input signal in deep convolutional networks get transformed into spikes in individual feature maps by learned filters. Summing and averaging over these spikes indicates the areas in the layers with most activity and provides an interpretable representation of the input/output. Similarly, the cABR signal summarizes peaks of neural activity as a response to the amplitude of the input stimulus.

The main advantage of these results is interpretability: the similarities in encoding are established by directly comparing individual acoustic features rather than performing linear transformations or correlation analyses. Our models are trained in a fully unsupervised manner in the GAN setting, where the Generator needs to learn to produce speech data not by replicating the input (as is the case in most models such as VAEs), but by imitation (producing data such that another network cannot distinguish it from real data). In the learning process, the networks generate innovative data^13^, which means the models feature one of the more prominent features of language—productivity.^77^ We test encoding of an acoustic property in networks that mimic both the production and perception principles and we test a phonetic property that encodes a phonological contrast (voiced vs. voiceless) in two languages.

Based on the common mechanism of averaging over neural activity in both signals, we can compare what other acoustic properties are encoded in second-to-last convolutional layers and in the brain stem. A detailed comparison of encoding of other acoustic properties is left for future work, but a test of which properties are encoded in both signals reveals several common properties. cABR signals have been shown to represent acoustic properties^73,74,78^ such as periodicity and the fundamental frequency (F0), lower frequency formants (e.g. F1, perhaps also F2;^79^), “acoustic onsets” such as burst, and “frequency transitions”^78^. Beguš and Zhou^50,51^ have shown that the same acoustic properties are encoded in the second to last convolutional layer as well based on a quantitative analysis of which acoustic properties are encoded in which convolutional layer. The following properties have been shown to be robustly encoded in the second to last convolutional layer: periodicity and F0 together with F0 transitions, low frequency formant structure (F1 and F2), burst, and timing of individual segments^50,51^. Figures 1; 3, 4; 6; and illustrate the similarities between the signal from intermediate convolutional layers (obtained by the proposed technique in Beguš and Zhou^50,51^) and the cABR signal. Later convolutional layers do not encode all these acoustic properties^50,51^. This suggests that many acoustic properties are encoded with frequency-following encoding only in the earlier layers of neural processing — both in the brain and in deep neural networks: F0, burst, timing, and low frequency formant structure. These parallels provide grounds for further explorations of how individual phonetic fearures are encoded in biological and artificial neural networks.

## 6 Conclusion and future directions

This paper presents a technique for comparing cABR neuroimaging of the brainstem with intermediate convolutional layers in deep neural networks. Both signals are based on summing and averaging of neural activity: either of electrical activity in the brain stem or of values in individual feature maps in convolutional layers. We argue that averaging over feature maps in deep convolutional networks parallels cABR recording in the brain because it summarizes areas in the convolutional layers with highest activity relative to the input/output. cABRs and second to last convolutional layers encode similar acoustic properties. Encoding of phonetic information is tested with cABR experiments on subjects of two different languages and with deep neural networks trained on these two languages. The results reveal that the two signals are highly similar and that encoding of phonetic features that result in phonological contrasts differ in similar ways in the brain stem and in intermediate convolutional layers between the two tested languages.

These results provide grounds for comparison of several other acoustic properties using the proposed framework, which are left for future work. Both intermediate convolutional layers and cABR signal represent several acoustic properties. World’s languages use various acoustic features to encode linguistically meaningful phonological contrasts. Testing these learned representations across different acoustic properties and languages should yield further information on similarities and differences in artificial and biological neural computation on speech data.

## Data availability

Data and checkpoints of trained models are available at: doi.org/10.17605/OSF.IO/ZDB52.

### Acknowledgements

Parts of this research were funded by a grant for new faculty at the University of California, Berkeley to G.B.

## Author contributions

**Gašper Beguš**: Conceptualization, Data curation, Formal Analysis, Investigation, Methodology, Project administration, Software, Supervision, Visualization, Writing – original draft; **Alan Zhou**: Conceptualization, Data curation, Formal Analysis, Investigation, Software, Writing – original draft; **T. Christina Zhao**: Conceptualization, Data curation, Resources, Writing – review & editing

## Footnotes

1 Norman-Haignere and McDermott^43^ propose a somewhat related experiment, where the stimulus is created such that it matches the natural stimulus in terms of model response.

2 Data can be accessed at https://osf.io/6fwxd/.

3 For slicing, we modified code written by Sameer Arshad for another study^12^.

4 Harwath and Glass^70^ propose a visualization technique for the DAVEnet model^71^ that involves summation — they operate with L2 norm values of individual filter activations, but they do not operate with the production (decoder) aspect of the networks and operate with spectrograms instead of waveforms. Their visualizations do not offer sufficiently high temporal resolution for comparison with the cABR signal (e.g. for vocalic periods). Their proposal additionally requires a PCA analysis for a comparison of intermediate convolutional layers with linguistically meaningful units. Their model does, however, show, that peak timing in intermediate convolutional layers correspond to segment boundaries (not vocalic peaks) in TIMIT.

